# Study of Basic Local Alignment Search Tool (BLAST) and Multiple Sequence Alignment (Clustal- X) of Monoclonal mice/human antibodies

**DOI:** 10.1101/2021.07.09.451785

**Authors:** Ivan Vito Ferrari, Paolo Patrizio

## Abstract

In this work, we have focused on the study of the Basic Local Alignment Search Tool (BLAST) and Multiple Sequence Alignment (Clustal-X) of different monoclonal mice antibodies to understand better the multiple alignments of sequences. Our strategy was to compare the light chains of multiple monoclonal antibodies to each other, calculating their identity percentage and in which amino acid portion. (See below figure 2) Subsequently, the same survey of heavy chains was carried out with the same methodology. (See below figure 3) Finally, sequence alignment between the light chain of one antibody and the heavy chain of another antibody was studied to understand what happens if chains are exchanged between antibodies. (See below figure 4) From our results of BLAST estimation alignment, we have reported that the Light Chains (Ls) of Monoclonal Antibodies in Comparison have a sequence Homology of about 60-80% and they have a part identical in sequence zone in range 100-210 residues amino acids, except ID PDB 4ISV, which it turns out to have a 40% lower homology than the others antibodies. As far as, the heavy chains (Hs) of Monoclonal Antibodies are concerned, however they tend to have a less homology of sequences, compared to lights chains consideration, equal to 60%-70% and they have an identical part in the sequence zone between 150-210 residues amino acids; with the exception of ID PDB 3I9G-3W9D antibodies that have an equal homology at 50%. (See supporting part) Summing up: about 70-80% identity among 2 light chains of 2 antibodies, 60-70% identity between 2 heavy chains of 2 antibodies, 30% identity between the two chains of a antibody and 30% if you compare the light chain of one antibody with the heavy chain of another antibody.

## 1. Introduction

- In bioinformatics, a sequence alignment is a way to organize DNA, RNA, or protein sequences to identify similarity regions that may be a consequence of functional, structural, or evolutionary relationships between sequences. Gaps are inserted between the residues so that identical or similar characters are aligned in successive columns [1-4]. It is important to consider that alignment makes it possible to identify identical or similar regions that may have functional, structural, or phylogenetic (evolutionary). Indeed, one the crucial aspect in this regard it is the sequence similarity which it is a powerful tool for discovering biological function. For to understand better what’s the biological function that we want to go and investigate, there are several online databases for multiple sequence alignments. One of the best known is the conserved Domain Database (CDD) that which is a compilation of multiple sequence alignments representing protein domains conserved in molecular evolution. NCBI’s Conserved Domain Database (CDD) is a resource for the annotation of protein sequences with the location of conserved domain footprints, and functional sites inferred from these footprints [5]. Depending on the type of sequence alignment (global or local) we find different software. The most used are :

‐ **BLAST** (Basic Local Alignment Search Tool): it finds regions of similarity between biological sequences. The program compares nucleotide or protein sequences to sequence databases and calculates the statistical significance [2,6].
‐ **Clustal**: Multiple Sequence Alignment: it used for Multiple alignments of nucleic acid and protein sequences. There are several software (Clustal Omega and ClustalW/ClustalX) [3,7]

In this work, we have focused on the study of the Basic Local Alignment Search Tool (BLAST) and Multiple Sequence Alignment (Clustal-X) of different Monoclonal mouse antibodies to better understand the multiple alignments of sequences.

## 2. Materials and Methods

### 2.1 BLAST (Basic Local Alignment Search Tool)

BLAST uses statistical theory to produce a bit score and expect value (E-value) for each alignment pair (query to hit). The bit score gives an indication of how good the alignment is; the higher the score, the better the alignment. It es statistical theory to produce a bit score and expect value (E-value) for each alignment pair (query to hit). The bit score gives an indication of how good the alignment is; the higher the score, the better the alignment [2,6]

#### 2.1.1 Parameters of BLAST

‐ Database (Non-redundant protein sequences)
‐ Organism (Mus musculus, taxed: 10090)
‐ Algorithm (blastp, protein-protein, BLAST)
‐ Max target sequences (100)
‐ Expect threshold (10)
‐ Word size (6)
‐ Matrix (BLOSUM 62)
‐ Gap costs (Existence: 11; Extension:1)

### 2.2. ClustalX (Multiple Sequence Alignment)

#### 2.21.1 Parameters of ClustalX

‐ Gap opening (10)
‐ Gap extension (0.2)
‐ Delay divergent sequences, % (30)
‐ DNA transition weight (0.5)
‐ Protein weight matrix (BLOSUM SERIES)

### 2.3 Protein Data Bank (https://www.rcsb.org/)

- **3W9D:** (Structure of Human Monoclonal Antibody E317 Fab)
- **1PSK:** (The Crystal structure of a Fab Fragment that binds to the melanoma-associated GD2-Ganglioside)
- **1F11:** (F124 Fab Fragment from a Monoclonal Anti-PRES2 Antibody)
- **1F58:** (IGG1 Fab Fragment (58.2) Complex with 24-residue peptide (Residues: 308-333 OF HIV-1 GP120 (MN isolate) with Ala to AIB substitution at position 323
- **1MF2:** (ANTI HIV1 Protease FAB Complex)
- **3O45:** (Crystal Structure of 101F Fab Bound to 17-mer Peptide Epitope)
- **1MPA:** (Bactericidal Antibody against Neisseria Meningitidis
- **3I9G:** (Crystal structure of the LT1009 (SONEPCIZUMAB) antibody Fab fragment in complex with sphingosine-1-phosphate)
- **3I50:** (Crystal structure of the West Nile Virus envelope glycoprotein in complex with the E53 antibody Fab)
- **3IET:** (Crystal Structure of 237mAb with antigen)
- **3VG9:** (Crystal structure of human adenosine A2A receptor with an allosteric inverse-agonist antibody at 2.7 A resolution)
- **4ISV:** (Crystal structure of the Fab Fragment of 1C2, a Monoclonal Antibody specific for Poly-Glutamine
- **4KVC:** (2H2 Fab fragment of immature Dengue virus)
- **4M43:** (Crystal structure of anti-rabies glycoprotein Fab 523-11)
- **5LQB:** (Complex structure of human IL2 mutant, Proleukin, with Fab fragment of NARA1 antibody)
- **6TYS:** (A potent cross-neutralizing antibody targeting the fusion glycoprotein inhibits Nipah virus and Hendra virus infection)

## 3. Results and Discussions

In simplistic terms antibodies perform two main functions in different regions of their structure. While one part of the antibody, the antigen binding fragment (Fab), recognizes the antigen, the other part of the antibody, known as the crystallizable fragment (Fc), interacts with other elements of the immune system, such as phagocytes or components of the complement pathway, to promote removal of the antigen. Antibodies all have the same basic structure consisting of two heavy and two light chains forming two Fab arms containing identical domains at either end attached by a flexible hinge region to the stem of the antibody, the Fc domain, giving the classical ‘Y’ shape. The chains fold into repeated immunoglobulin folds consisting of anti-parallel β-sheets (1), which form either constant or variable domains. The Fab domains consist of two variable and two constant domains, with the two variable domains making up the variable fragment (Fv), which provides the antigen specificity of the antibody, with the constant domains acting as a structural framework. Each variable domain contains three hypervariable loops, known as complementarity determining regions (CDRs), evenly distributed between four less variable framework (FR) regions. It is the CDRs that provide a specific antigen recognition site on the surface of the antibody and the hypervariability of these regions enables antibodies to recognize an almost unlimited number of antigens. [8] (See below figure 1) The aim of this work was to fully understand, what are the potentials of sequence alignment, estimated by BLAST and ClustalX method. Our strategy was to compare the light chains of multiple monoclonal antibodies to each other, calculating their identity percentage and in which amino acid portion. (See below figure 2) Subsequently, the same survey of heavy chains was carried out with the same methodology. (See below figure 3) Finally, sequence alignment between the light chain of one antibody and the heavy chain of another antibody was studied to understand what happens if chains are exchanged between antibodies. (See below figure 4) From our results of BLAST estimation alignment, we have reported that the Light Chains (Ls) of Monoclonal Antibodies in Comparison have a sequence Homology of about 60-80% and they have a part identical in sequence zone in range 100-210 residues amino acids, except ID PDB 4ISV, which it turns out to have a 40% lower homology than the others antibodies. As far as, the heavy chains (Hs) of Monoclonal Antibodies are concerned, however they tend to have a less homology of sequences, compared to lights chains consideration, equal to 60%-70% and they have an identical part in the sequence zone between 150-210 residues amino acids; with the exception of ID PDB 3I9G-3W9D antibodies that have an equal homology at 50%. (See supporting part)

**Fig 1.**
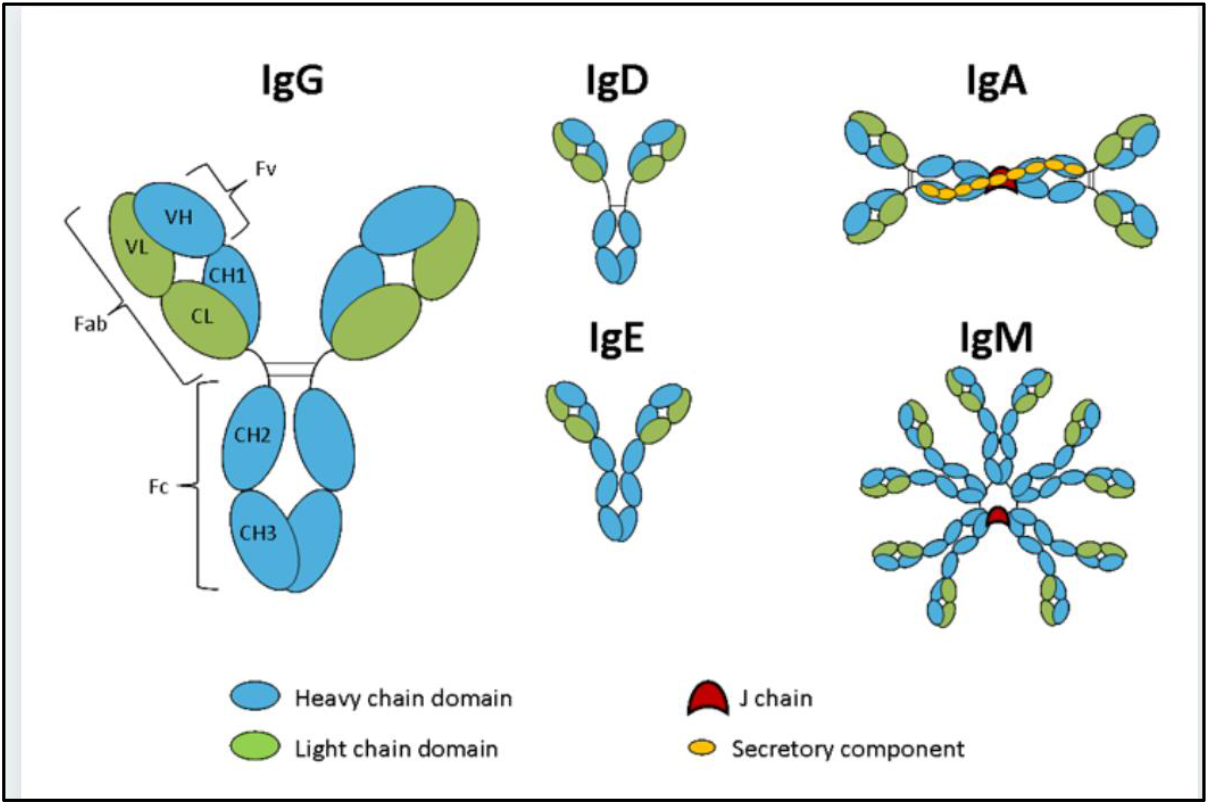
Schematic representation of the five immunoglobulin classes or isotypes in mammals. Figure reproduce from https://absoluteantibody.com/antibody-resources/antibody-overview/antibody-isotypes-subtypes/

**Fig 2.**
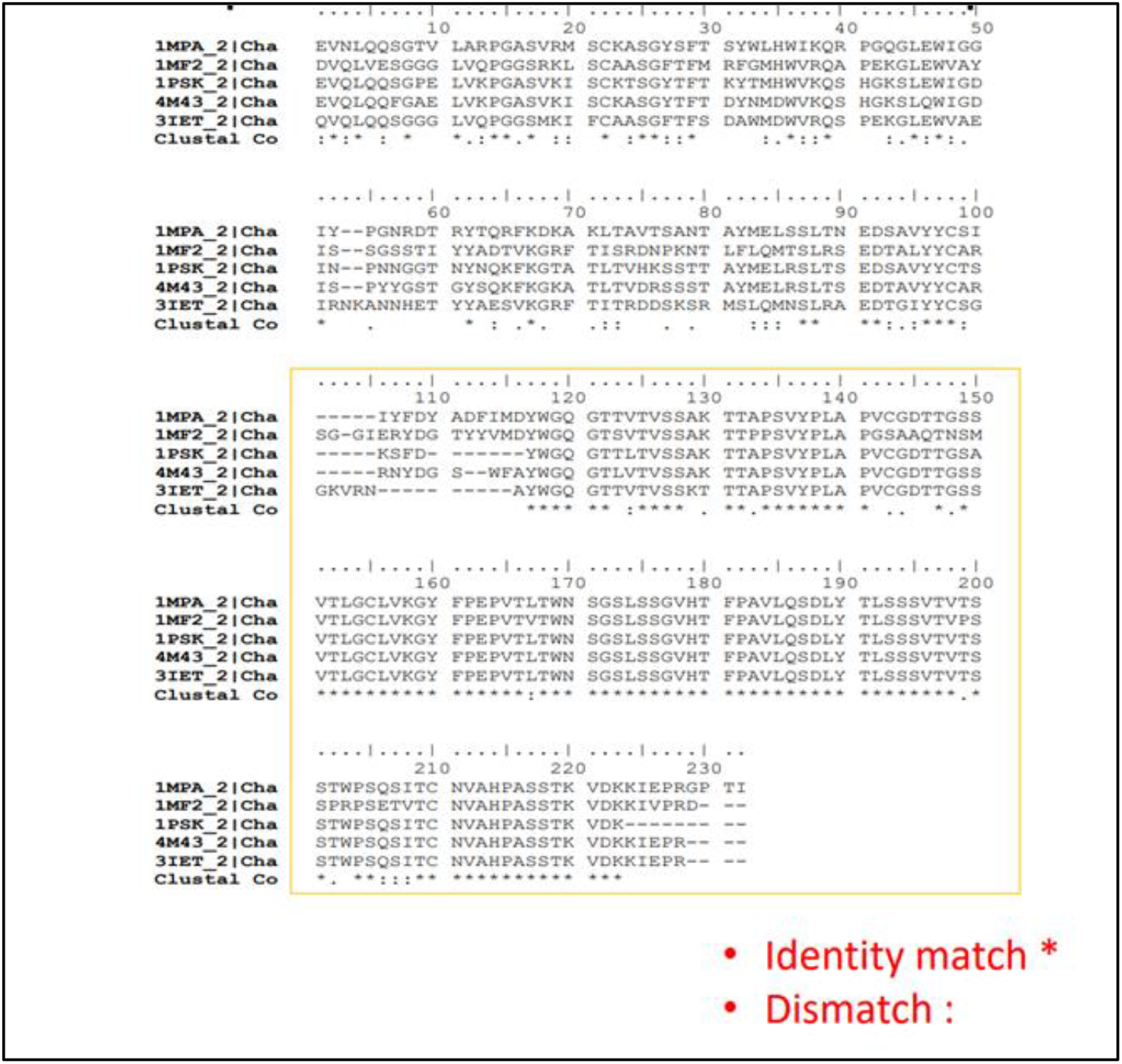
Comparison of two light chains between 5 antibodies

**Fig 3.**
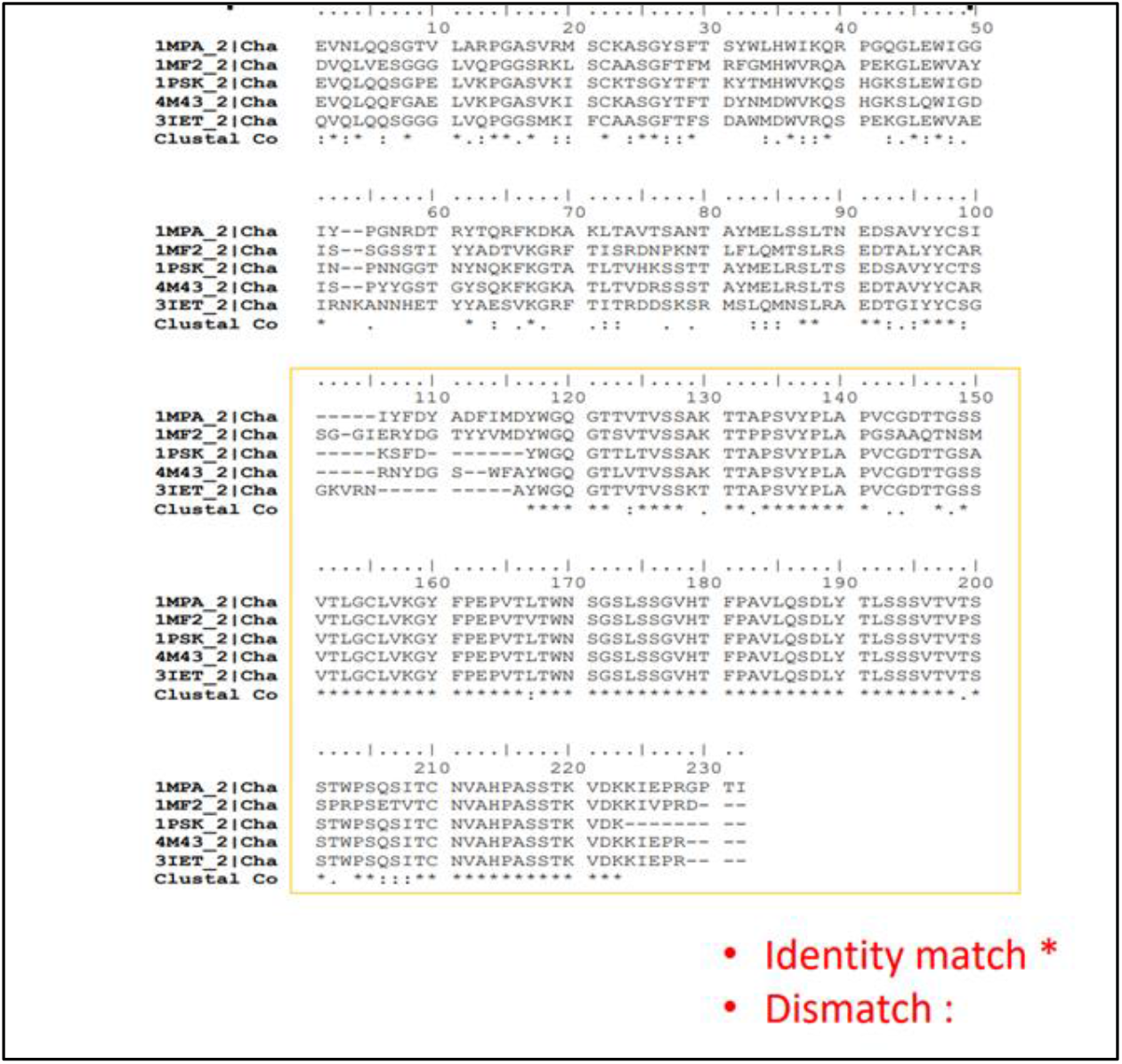
Comparison of two heavy chains between 5 antibodies

**Fig 4.**
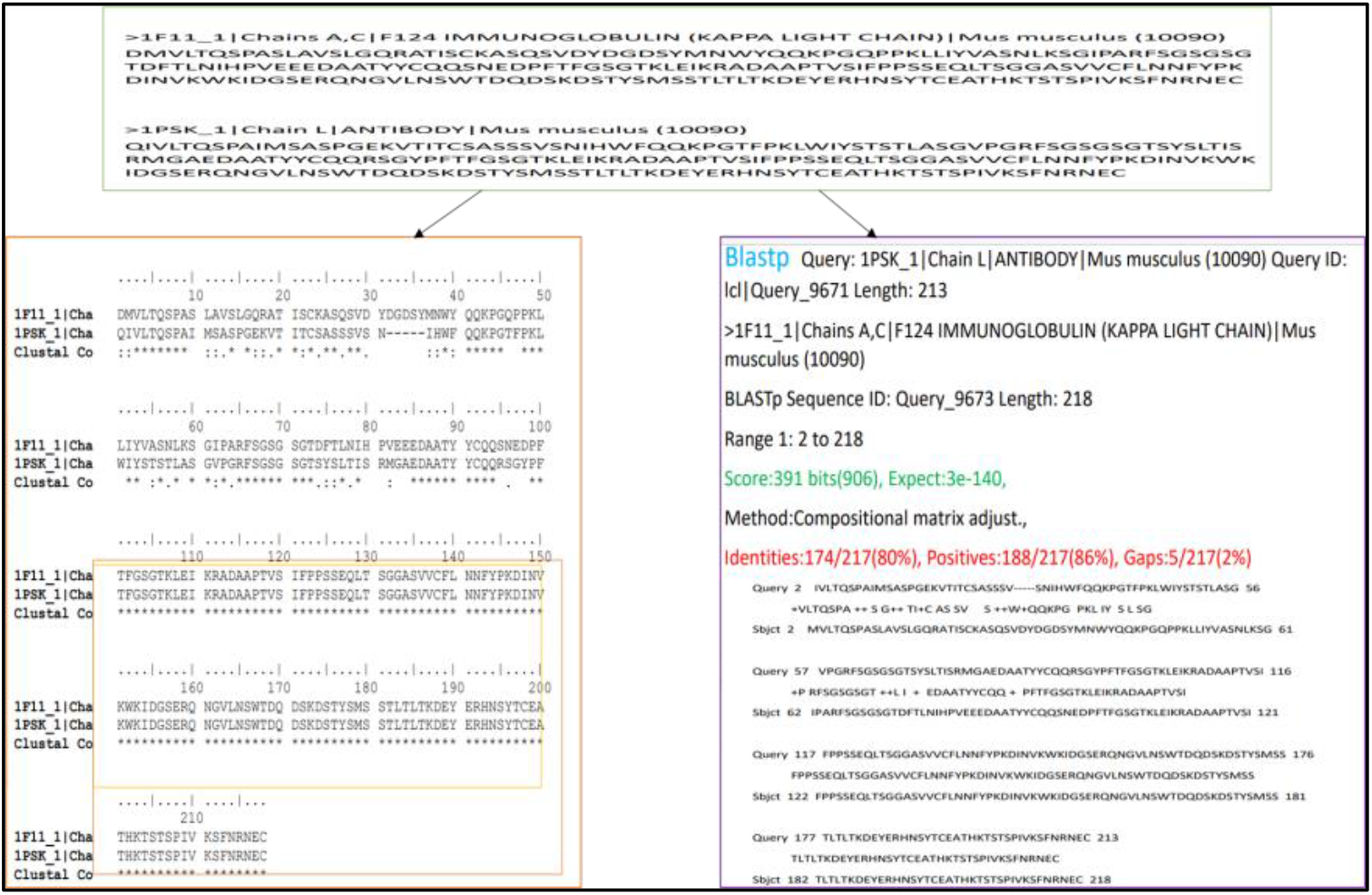
Comparison of two light chains (ID PDB 1PSK and ID PDB 1F11) of antibodies, estimated on the left side by ClustalX and on the right side by BLAST

To understand better this type of investigation, we have examined two examples of monoclonal antibodies, first comparing their light chains to each other and then their heavy chains, estimating with BLAST their percentage of identity and similarity with each other. (See below figure 4-5;6-7) As can be seen from Figure 4-7, we found that the identity percentage of the 2 light chains/ heavy chains of the 2 antibodies compared is about 80%, while the similarity is about 86%, the 2% gaps. The last survey we want to present in this study is to try to understand by asking a question: what would happen if sequence alignments are made between the light chain of an antibody with the heavy chain of another antibody? Would the percentage of identity and similarity of sequence increase or decrease, relative to their own? To get to the bottom of the matter we investigated 11 monoclonal antibodies by making all the combinations between the light chain of an antibody with the heavy chain of another antibody. (See supporting part and See below figure 8)

**Fig 5.**
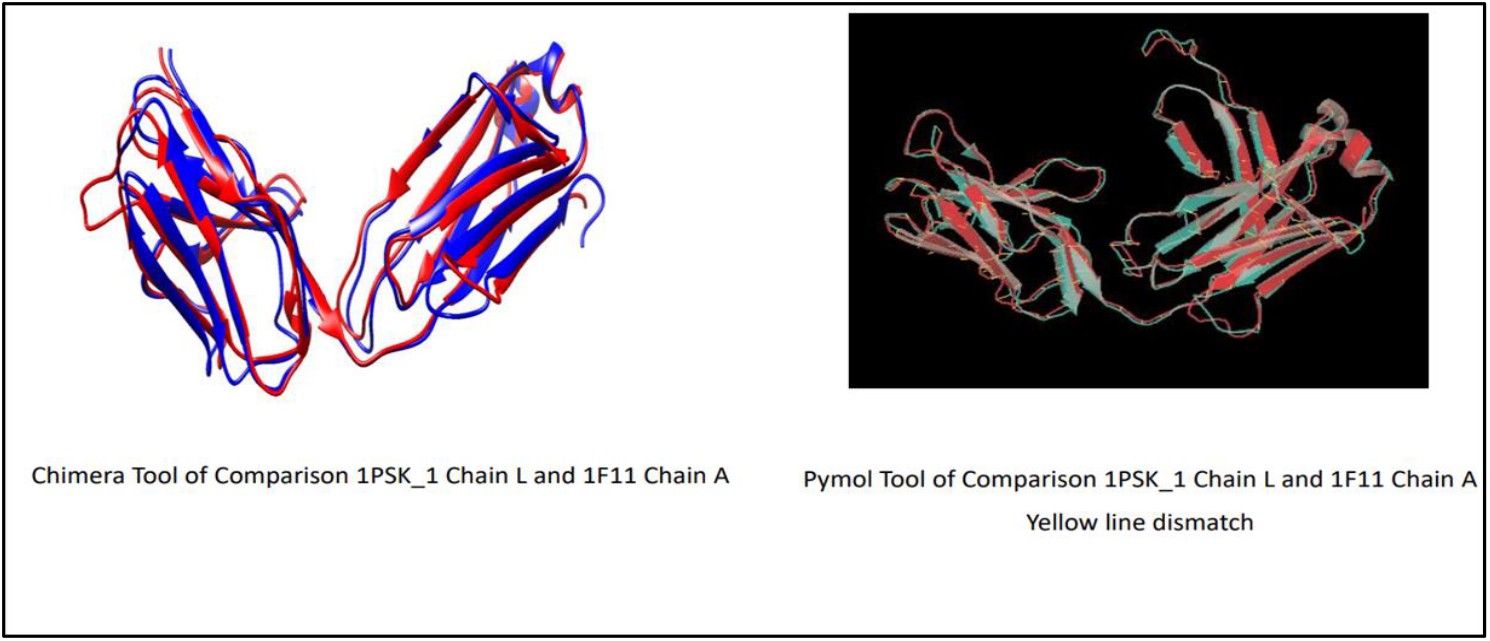
Comparison of two light chains (ID PDB 1PSK and ID PDB 1F11) of antibodies. Figure reproduced, on the left side by Chimera Software and on the right side by Pymol Software

**Fig 6.**
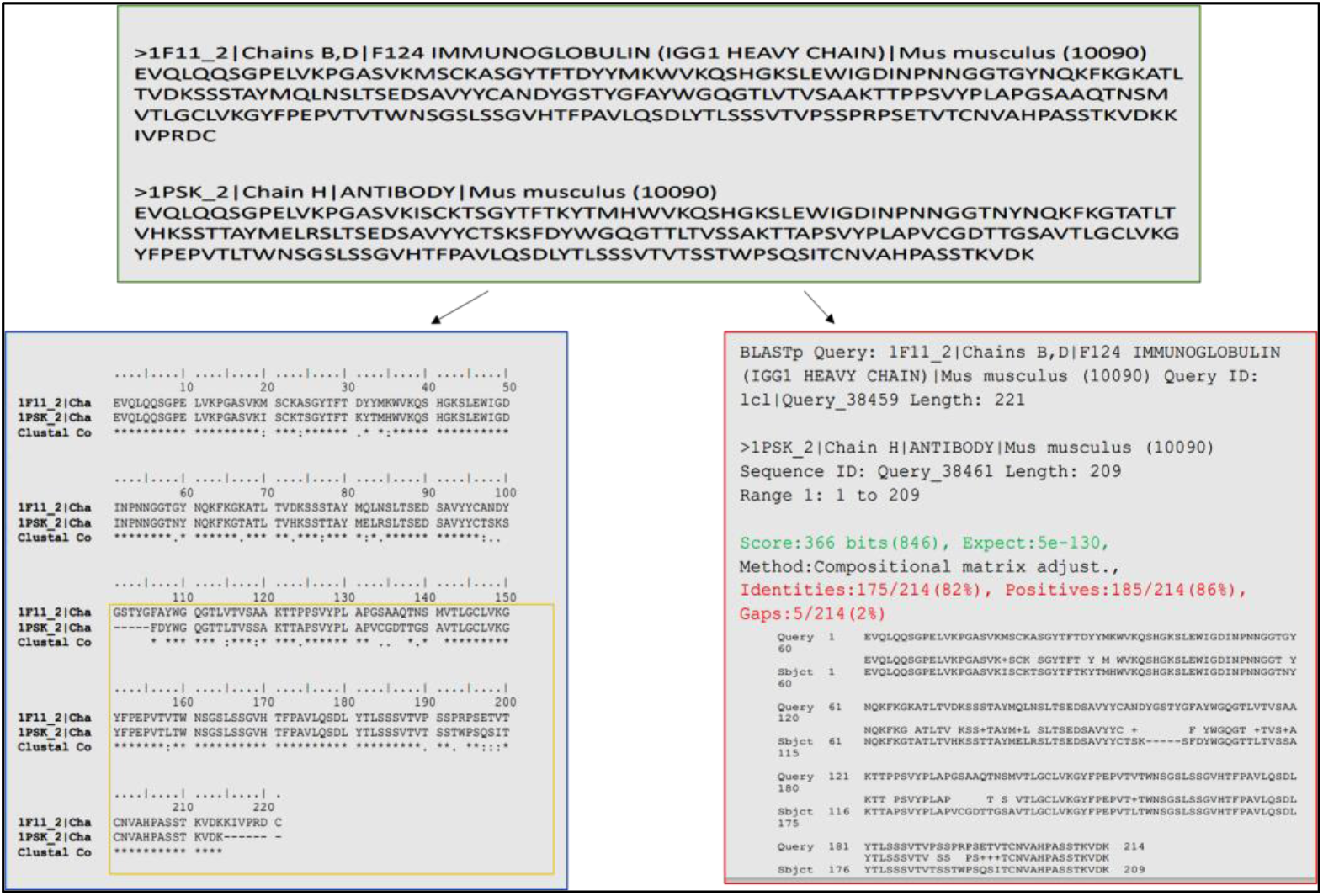
Comparison of two heavy chains (ID PDB 1PSK and ID PDB 1F11) of antibodies, estimated on the left side by ClustalX and on the right side by

**Fig 7.**
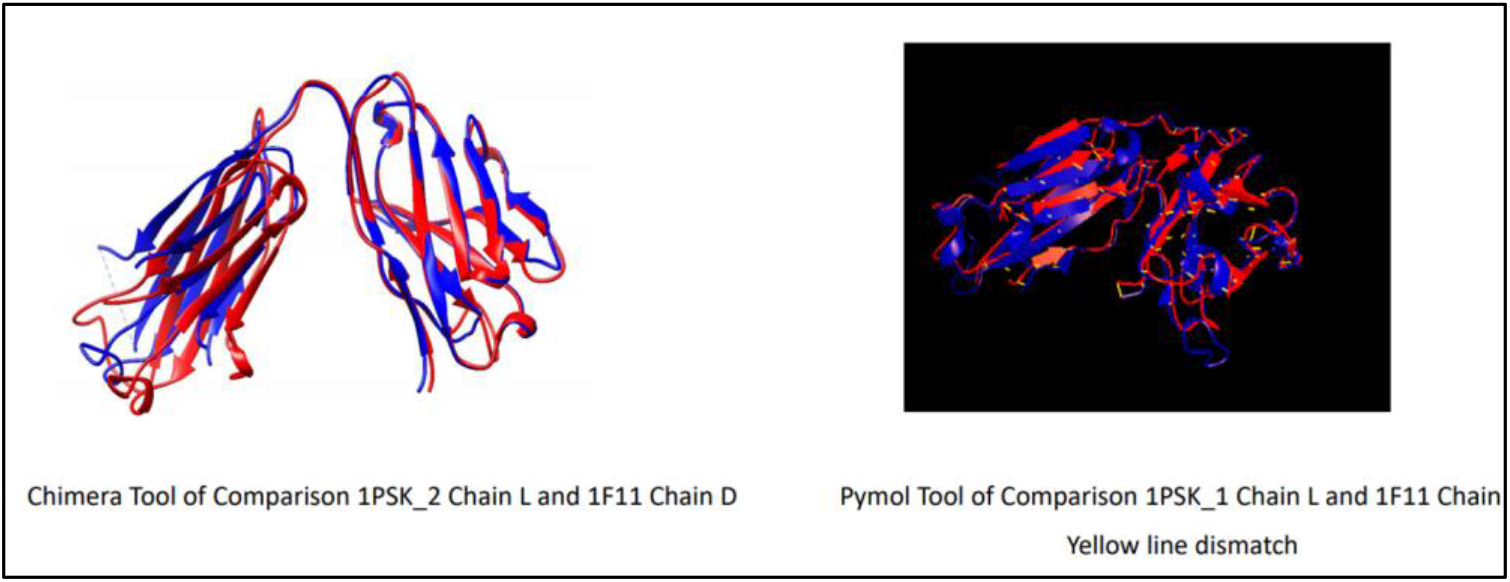
Comparison of two heavy chains (ID PDB 1PSK and ID PDB 1F11) of antibodies. Figure reproduced, on the left side by Chimera Software and on the right side by Pymol Software

**Fig 8.**
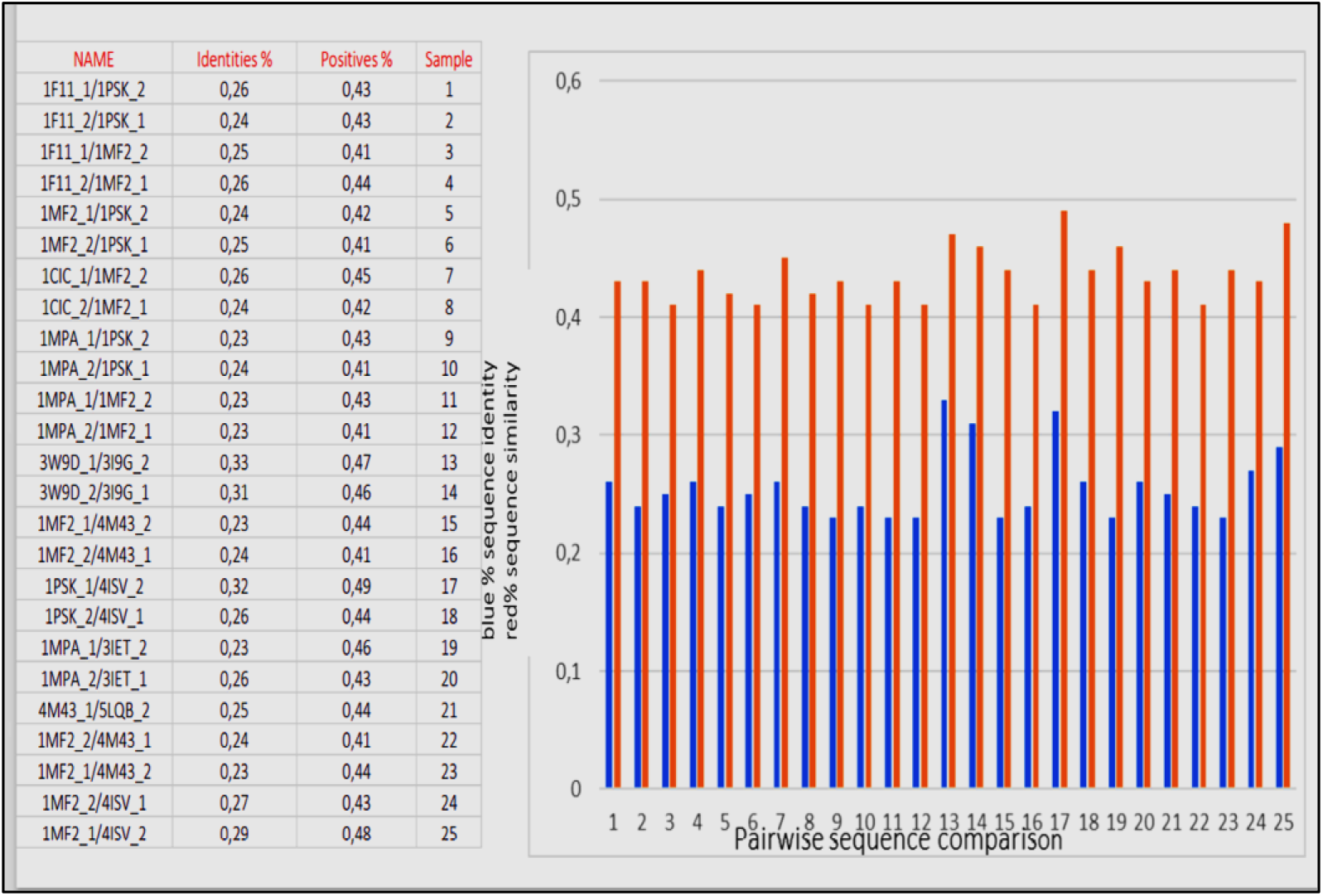
Comparison between the light chain of an antibody and heavy chain of another antibody and vice versa, estimated by BLAST method

## 4. Conclusion

In this work, we have focused on the study of the Basic Local Alignment Search Tool (BLAST) and Multiple Sequence Alignment (Clustal-X) of different Monoclonal mouse antibodies to better understand the multiple alignments of sequences. Our strategy was to compare the light chains of multiple monoclonal antibodies to each other, calculating their identity percentage and in which amino acid portion. (See below figure 2) Subsequently, the same survey of heavy chains was carried out with the same methodology. (See below figure 3) Finally, sequence alignment between the light chain of one antibody and the heavy chain of another antibody was studied to understand what happens if chains are exchanged between antibodies. (See below figure 4) From our results of BLAST estimation alignment, we have reported that the Light Chains (Ls) of Monoclonal Antibodies in Comparison have a sequence Homology of about 60-80% and they have a part identical in sequence zone in range 100-210 residues amino acids, except ID PDB 4ISV, which it turns out to have a 40% lower homology than the others antibodies. As far as, the heavy chains (Hs) of Monoclonal Antibodies are concerned, however they tend to have a less homology of sequences, compared to lights chains consideration, equal to 60%-70% and they have an identical part in the sequence zone between 150-210 residues amino acids; with the exception of ID PDB 3I9G-3W9D antibodies that have an equal homology at 50%. (See supporting part) Summing up: about 70-80% identity among 2 light chains of 2 antibodies, 60-70% identity between 2 heavy chains of 2 antibodies, 30% identity between the two chains of a antibody and 30% if you compare the light chain of one antibody with the heavy chain of another antibody.

## Conflicts of Interest

The authors declare no conflict of interest

## Notes

### Competing Interest Statement

The authors have declared no competing interest.

